# Fronto-parietal, cingulo-opercular and striatal contributions to learning and implementing control policies

**DOI:** 10.1101/2020.05.10.086587

**Authors:** Apoorva Bhandari, David Badre

## Abstract

Efficient task performance requires co-ordination of internal cognitive processes by implementing control policies adapted to the dynamic structure of task demands. The cognitive and neural basis of control policy implementation remains poorly characterized, in part because it is typically confounded with implementing new stimulus-response rules. To disambiguate these processes, we asked participants to perform multiple novel variants of a working memory control task. Each variant had a unique, novel sequential trial structure, but all shared common stimulus-response rules, enabling us to test control policy implementation separate from rule learning. Behaviorally, we found evidence for two adaptive processes tied to control policy implementation. One process was reflected in slower responses on the first trial with a novel sequential trial structure, followed by rapid speeding on subsequent trials. A second process was reflected in the diminishing size of the first trial cost as participants accommodated different variants of the task over many blocks. Using fMRI, we observed that the striatum and a cingulo-opercular cortical network increased activity to the first trial, tracking the fast adjustment. This pattern of activity dissociated these regions from a fronto-parietal network including dorsolateral PFC, inferior frontal junction, inferior parietal sulcus, and rostrolateral PFC, which showed a slower decline in activity across trials, mirroring findings in rule implementation studies, but in the absence of rule implementation demands. Our results reveal two adaptive processes underlying the implementation of efficient, generalizable control policies, and suggest a broader account of the role of a cortico-striatal network in control policy implementation.

**Significance statement:** Rapid adaptation to novel tasks is a hallmark of human behavior. Understanding how human brains achieve this is of critical importance in neuroscience. Here we broaden the scope of this problem, going beyond task rules to more broadly consider the cognitive control demands produced by novel task dynamics. We propose that humans rely on two adaptive processes to rapidly implement efficient, generalizable control policies as task dynamics change, even when task rules remain unchanged. One process unfolds rapidly and underlies efficient adaptation. A second process unfolds slowly with experience across task conditions and underlies generalization of control policies. Using fMRI, we identify cingulo-opercular cortex, fronto-parietal cortex and striatum as dissociable components of a cortico-striatal network that contribute to control implementation.

## Introduction

Everyday tasks place complex and dynamic demands on internal cognitive processes like perception, attention, memory retrieval and action selection. When a task is novel, the moment-to-moment co-ordination of these processes requires *cognitive control* (Miller and Cohen, 2001). As a result, we perform new tasks slowly and with errors, at least initially (Schneider and Shiffrin, 1977; Norman and Shallice, 1986). But with experience, we quickly settle on a cognitive routine for efficient task performance (Cole et al., 2010; Ruge and Wolfensteller, 2010; Wolfensteller and Ruge, 2012; Bhandari and Duncan, 2014; Bhandari and Badre, 2018). Such rapid ‘task automatization’ (Mohr et al., 2016) is observed across a wide variety of tasks. This suggests that we can rapidly and flexibly implement an *internal control policy* that enables efficient coordination of cognitive processing (Bhandari and Duncan, 2014; Bhandari and Badre, 2018; Chiu and Egner, 2019). The neural systems that implement control policies remain understudied.

Previous studies of task automatization have primarily focused on learning and implementation of task rules (stimulus-response contingencies) through trial-and-error (Cao et al., 2016) or instruction (Brass and Von Cramon, 2002; Brass and Cramon, 2004; Cole et al., 2010; Ruge and Wolfensteller, 2010; Dumontheil et al., 2011; Hartstra et al., 2011; Stocco et al., 2012; Mohr et al., 2016; Muhle-Karbe et al., 2017; Bourguignon et al., 2018; Hampshire et al., 2019; Ruge et al., 2019). These studies have implicated the fronto-parietal networks and striatum in automatization. In particular, several studies have reported declining activity in fronto-parietal networks correlated with diminishing response time (RT) over practice or following the first trials of a task (Cole et al., 2010; Ruge and Wolfensteller, 2010; Hartstra et al., 2011; Muhle-Karbe et al., 2017; Hampshire et al., 2019). Mohr et al. (2016) observed enhanced coupling between dorsal attention and cingulo-opercular networks with practice. Subcortically, Ruge & Wolfensteller (2013, 2015) found higher activation in striatum and greater fronto-striatal coupling with practice.

However, rules are only one type of knowledge required for task implementation. Tasks also possess dynamic structure in the order and timing of events (Radvansky & Zacks, 2014). Such structure affects which mental operations are relevant at any moment. Thus, efficient task performance requires control policies aligned to its dynamic structure.

As an example, consider the task in Figure 1. Three cues are presented in a sequence. One cue provides a context (top-level) that determines which of the other two (bottom-level) cues is a target or distractor based on known rules. The target cue must be maintained in working memory for a subsequent decision.

**Figure 1.**
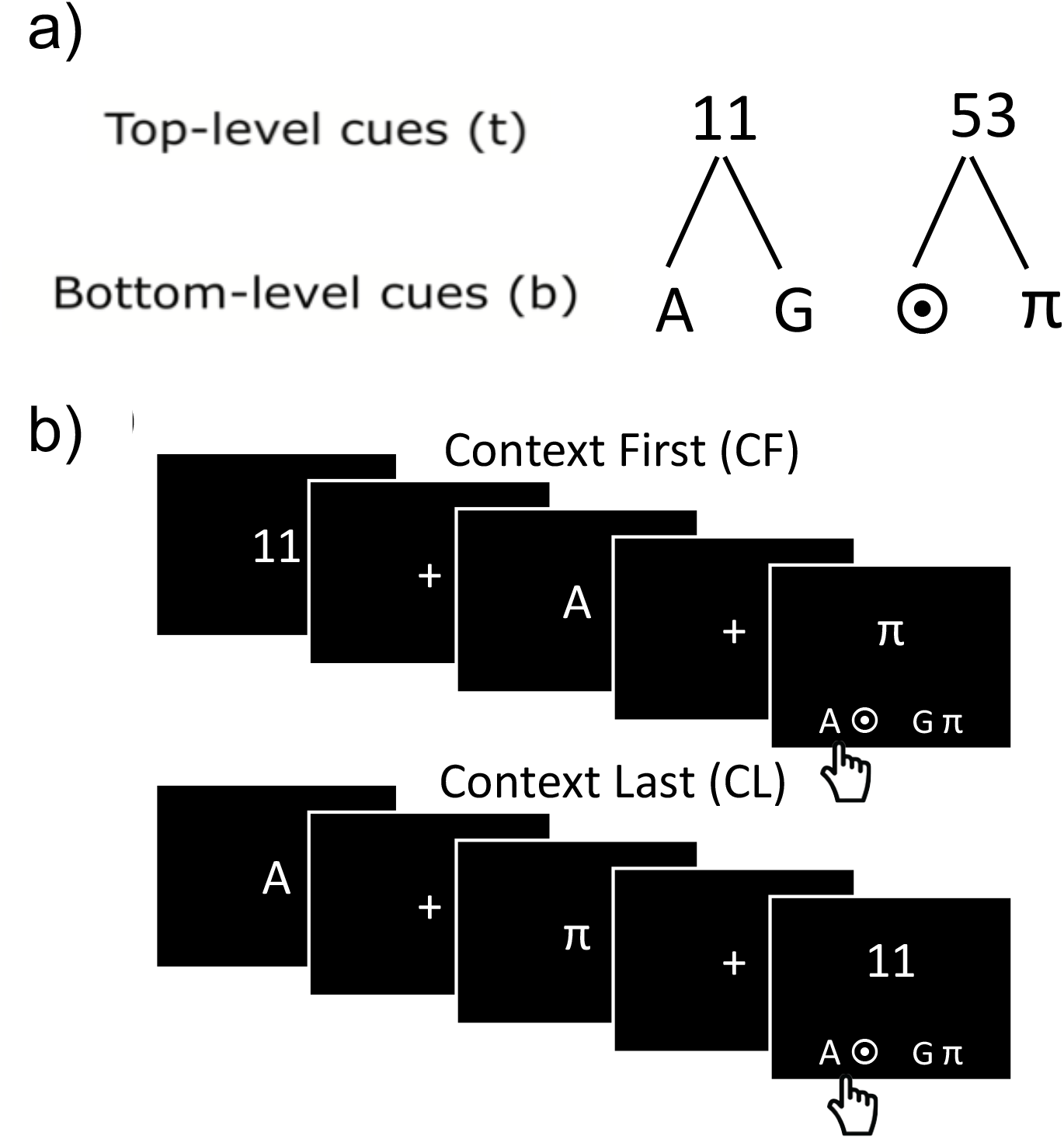
Working memory control task employed by Bhandari & Badre (2018). (a) Rule trees linking top-level (contextual) cues bottom-level cues. Participants are asked to memorize the rule trees prior to beginning the task. (b) Sample context first (CF) and context last (CL) trials. The top-level cue (number) indicated which of the other two cues in the sequence is the target. Each trial concluded with a response panel that consisted of two pairs of items. Participants indicated with a key press which pair contained the correct target.

The policy for this task depends on the cue sequence. On ‘context last’ (CL) trials, the toplevel cue appears last. Thus, participants must store both bottom-level cues in working memory, and then selectively ‘output gate’ the target item once the context appears (‘output-gating policy’). On ‘context first’ (CF) trials, while the same policy could be used, a more efficient one is available. The context can guide selectively input-gating only the target bottom-level item in working memory (‘input-gating policy’). Therefore, different sequential structures afford distinct control policies.

In a recent study, we gave one group of participants a pure block of CL trials, followed by a pure CF block, while another group received two CF blocks. Participants in the first group learned an output-gating policy to solve the task. They then transferred this policy to the CF block, evident in slower initial RT compared to participants who only did two CF blocks. Moreover, this transfer from CL to CF occurred even when new stimulus-response rules were instructed before the second CF block. With experience, participants transitioned to the more efficient input-gating policy.

Thus, people implement control policies aligned to tasks’ dynamic structures, and transfer them to novel contexts independent of task rules. A key open question, however, is what neural systems support control policy learning and implementation.

## Methods

### Overview of the current experiment

In the current study, we examined the neural substrates of control policy implementation, disentangling them from rule learning. We hypothesized that multiple-demand regions associated with cognitive control in the frontal and parietal cortex, as well as the striatum would support the implementation of control policies, even when demands to learn new stimulus-response relationships are minimal. To test this hypothesis, we developed a 3^rd^ order version of the working memory control task (Figure 2), which afforded a larger space of possible sequential trial structures. From this space, we generated twenty unique sequential trial structures, but all sharing identical stimulus-response rules. This enabled us to repeatedly measure brain activity as participants implemented control policies aligned to novel sequence structures, and while the task rules remained unchanged.

**Figure 2.**
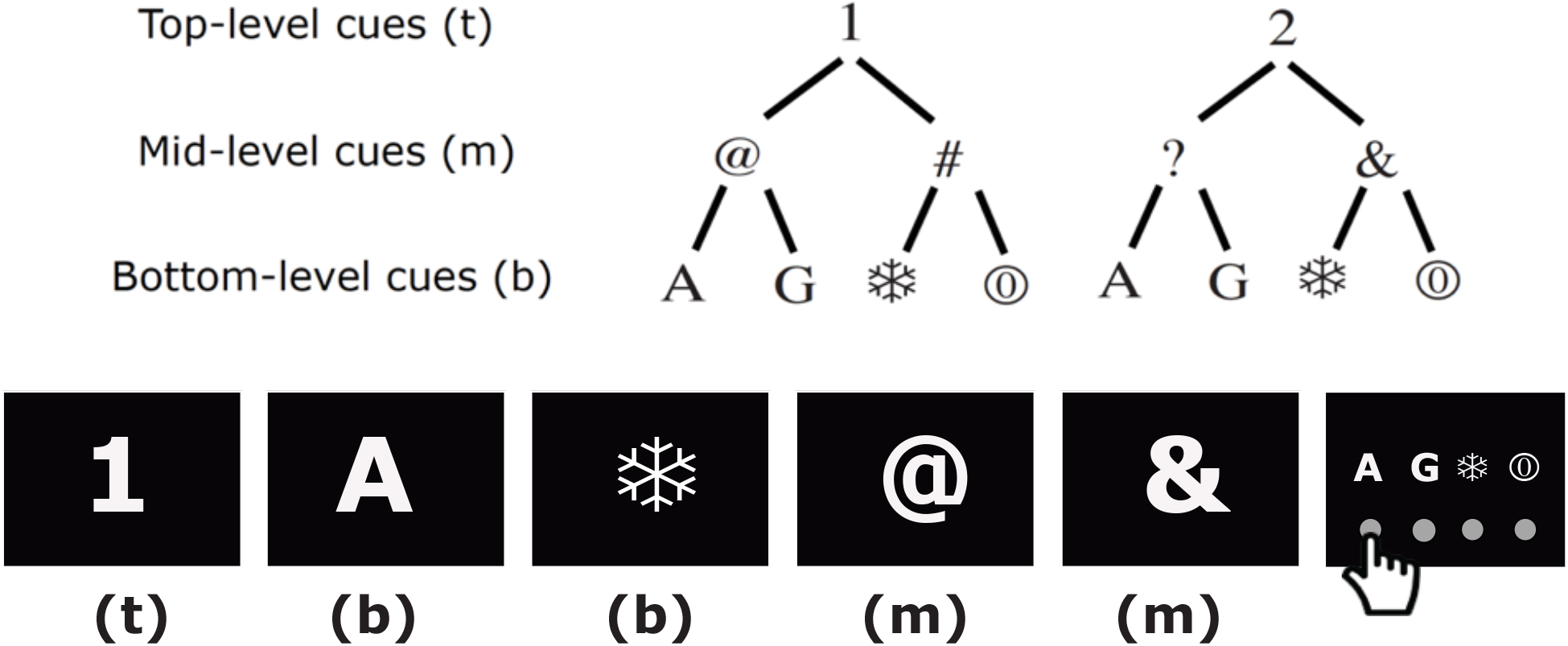
Experimental task. Participants were trained on a 3^rd^ order working memory control task. Top panel shows the ‘rule trees’ that specified the hierarchical relationships between top, mid and bottom level cues. Participants were extensively trained to memorize these rules, which remained unchanged throughout the experiment. Bottom panel shows a sample trial. Each trial consisted of a 5-item sequence containing one top-level, two midlevels and two bottom-level cues, and a response panel which was presented at the same time as the 5^th^ item. On every trial, the hierarchical rules uniquely specified a bottom-level cue that served as the target. Participants were asked to identify this target bottom-level cue and indicate their choice by pressing a button corresponding to the position of the chosen target in the response panel. In the sample trial shown, the sequential trial structure [(t)-(b)-(b)-(m)-(m) for the sample trial] constrains this step-wise process of identifying the target ‘A’ and thus determines the moment-by-moment control demands of the task. A novel sequential trial structure is employed in every block.

We predicted that control policy implementation would be indexed by the rapid speeding of responses over the first few trials of experience with a given sequential trial structure and correlates of such behavioral dynamics would be reflected in the activity dynamics and functional coupling of regions in cortico-striatal networks, in particular task control networks.

### Participants

26 adult, right-handed participants (13 males, 13 females; age-range: 18-29, M = 22.5, SD = 3) were recruited from the Providence, RI area. All participants gave informed, written consent as approved by the Human Research Protections Office of Brown University, and they were compensated at the rate of $20/hour for their participation. A total of 5 participants were excluded from the study. 2 participants were excluded due to very low performance (< 60%) and indication of non-compliance with task instructions. 1 participant was excluded due to their vision being obstructed by condensation on MRI-safe glasses during scanning, and the resulting disruption to task performance. 2 participants were excluded due to excessive head movement (> 3mm) during MRI scanning. Thus, fMRI data from 21 participants (10 males, 11 females; age-range: 18-29, M = 22.4, SD = 3) advanced to analysis. All participants had normal or corrected-to-normal vision, and no reported neurological or psychological disorders.

### Experiment Design & Statistical Analysis

#### Behavioral Task

Participants were instructed to perform the 3^rd^ order working memory control task shown in Figure 2. They were instructed to observe a sequence of five visual cues presented on the screen and to identify, remember and subsequently report the one target cue among these five. The cue sequence was constructed from the ‘rule trees’ shown in the top panel of Figure 2. Each sequence had one top-level cue (either 1 or 2), one middle-level item from each tree (@ or #, and, ? or &), and two lower-level items – a letter (either A or G) and a symbol (either ❄ or ☉). The top-level item served as a contextual cue for specifying which of the two middle-level cues was relevant on the trial. The relevant middle-level cue, in turn, specified which of the two lower level cues – letter or symbol – was the target. At the end of the sequence, coincident with the final cue, participants were presented with a four-item response panel that consisted of all the four possible bottom-level cues. They indicate with a four-alternative button press which cue they identified as the target. The order (left to right) of the four cues in the response panel, and so their assignment to a button press response, was randomized on a trial-by-trial basis.

Each trial of the task may be construed as a search through the rule trees for a branch that includes at least one presented top-level, middle-level and lower-level item (Figure 2(b)). Thus, the task is necessarily solved in steps as stimuli are revealed in sequence. Crucially, the *order* in which the top-level, middle-level and lower-level cues are presented determines the moment-by-moment control demands of the task. For example, if the top-level cue is presented prior to the middle-level cues, then the former can be used to selectively store the relevant middle-level cue in working memory. On the other hand, if the middle-level cues are presented first, then both must be stored in working memory, and the top-level cue is subsequently employed to selectively retrieve the relevant middle-level item. Thus, the sequential trial structure is a task grammar that can be represented in abstract form by identifying the order in which the top, bottom and middle level cues appear. For example, the trial shown in Figure 2(B) is an instance of the sequential trial structure (t)-(b)-(b)-(m)-(m). We assume that each sequential trial structure produces distinct control demands and so, given a block of trials generated from a novel sequential trial structure, participants would need to learn or adapt a control policy to the specific demands of that structure.

Cues in the sequence were each presented for 500 ms with a fixed stimulus onset asynchrony (SOA) of 1s. A fixation cross was displayed between the termination of one cue and the onset of the next. The fifth cue was presented at the same time as the response panel and terminated at 500 ms or when a response was made, whichever was earlier. The response panel continued to be on the screen until a response was made. Participants were given up to 5 seconds to respond, but were asked to respond as quickly as possible while minimizing errors.

#### Experimental protocol

Experimental sessions began with a three-part training protocol administered to all participants to aid them in learning the rules of the task.

First, participants were instructed on a simpler task for which the rules consisted of the top half of the two rule trees (i.e. only including top and middle-level cues). Each trial consisted of 3-item stimulus sequences: a single top-level item and one middle-level item from each of the two trees. Participants were asked to remember and subsequently report the target cue; i.e., the middle-level cue associated with the top-level cue. After each response, they received auditory feedback (Correct responses: high tone [400 Hz] for 200 ms; Incorrect or missed responses: low tone [200 Hz] for 200 ms). Participants performed this task until they completed a full block of 16 trials with 1 error or less.

Second, participants were instructed on another simpler task that was formed out of the bottom half of the two rule trees (i.e. the four smaller trees formed when including only middle-level and bottom-level cues). Again, each trial consisted of 3-item stimulus sequences - a single middle-level item, and one bottom-level cue of each type (i.e. letters and symbols). Participants were asked to remember and subsequently report the target cue - i.e. the bottom-level cue associated with the middle-level cue. Participants received visual and auditory feedback. Again, participants performed this task until they performed a full block of 16 trials with 1 error or less.

Third, participants were instructed on the full task. Each trial now consisted of 5-item stimulus sequences as described earlier. For training, a single sequential trial structure, (m)-(b)-(b)-(t)-(m), was employed. Again, participants performed this task until they performed a full block of 16 trials with 1 error or less. Finally, participants also performed one block of 16 trials without feedback to acclimatize them to performing without feedback, as would be the case in the main experiment. Note that previous work has indicated that piecewise training of hierarchical rules does not impose a particular order of sequential decisions (Ranti et al., 2016).

In order to adequately sample the control policy learning process, we generated trials from 20 distinct sequential trial structures. These trial structures are listed in Table 1. Trials were organized into 20 blocks of 12 trials, with each block consisting of trials generated from a single, novel sequential trial structure. Importantly, all 20 structures employed in the testing session were distinct from the one employed during training and the task rules were unchanged. Therefore, trained participants were highly familiar with the task rules, but were naïve to all the sequential trial structures presented in the test session. At the beginning of each block, the task rules were presented on the screen for 8 s followed by 8s of fixation. The 12 trials were then presented. The inter-trial fixation interval was jittered between 1 and 12 seconds, with a mean ITI of 4.3 seconds, optimized for maximizing the efficiency of deconvolving the response to each trial in the block from the one before and after it. Participants were also given 8 s of fixation between two blocks. Additionally, after the completion of ten blocks, participants were given a 90 s break during which time they were asked to rest their eyes while staying still.

**Table 1:** Sequential trial structures employed

| **Training:** (t)-(b)-(b)-(m)-(m) |  |
| --- | --- |
| **Testing** |  |
| (m)-(b)-(b)-(t)-(m) | (m)-(b)-(m)-(b)-(t) |
| (b)-(t)-(m)-(m)-(b) | (b)-(m)-(b)-(m)-(t) |
| (m)-(b)-(b)-(t)-(m) | (b)-(t)-(b)-(m)-(m) |
| (m)-(b)-(t)-(b)-(m) | (b)-(t)-(m)-(b)-(m) |
| (m)-(m)-(b)-(b)-(t) | (b)-(b)-(m)-(t)-(m) |
| (m)-(t)-(b)-(b)-(m) | (b)-(m)-(m)-(t)-(b) |
| (t)-(m)-(b)-(m)-(b) | (m)-(t)-(b)-(m)-(b) |
| (b)-(b)-(m)-(m)-(t) | (t)-(b)-(m)-(b)-(m) |
| (b)-(m)-(t)-(m)-(b) | (m)-(b)-(b)-(t)-(m) |
| (m)-(b)-(t)-(b)-(m) | (b)-(b)-(t)-(m)-(m) |

#### Behavioral Analysis

Control policy implementation was behaviorally indexed by changes error rates and response times as a function of trial experience with a novel sequential trial structure. These changes were statistically evaluated with one-way, repeated measure ANOVAs. Post-hoc, pairwise comparisons between levels of trial experience were evaluated with paired *t*-tests and Bonferroni-corrected for multiple comparisons. To evaluate changes in rates of responses time speeding across, exponential decay functions were fit to the mean response time by trial number curves and decay parameters were compared with paired *t*-tests.

#### fMRI Procedure and Preprocessing

Whole-brain imaging was performed on a Siemens 3T PRISMA MRI system equipped with a 32-channel head coil. A high-resolution, T1-weighted 3D multiecho MPRAGE anatomical image was obtained from each participant for the purposes of visualization. Functional data were acquired in a single run of ~1800 volumes using a gradient-echo, echoplanar pulse sequence (TR 2 s, TE 28 ms, flip angle 90 deg, 38 interleaved axial slices, 192 mm field of view, 3 x 3 x 3 mm voxel size). Stimuli were presented on a display screen located behind the scanner and made visible to the participant via an angled mirror attached to the head coil. Cushioned padding was placed around the head to restrict head motion. Participants made their responses with their right hand using an MRI-compatible button box.

#### fMRI Analyses

Preprocessed fMRI data were analysed with a statistical parametric mapping approach in SPM12. A general linear model was constructed with 2 sets of 12 trial epoch regressors, one set for each trial position for the first 10 of the 20 blocks of the experiment (set 1) and another for the last 10 blocks (set 2). Each trial regressor was specified as a set of box-cars covering the period from the onset of the first cue to the time of the onset of the final cue at the end of the trial for all relevant trials. Error-trials (regardless of trial position within the block) were modelled separately. Trial-by-trial RTs were separately modelled. In addition, we included 4 separate key-press stick regressors to model motor-related activity, as well as estimates of translational and rotational motion as covariates of no-interest. In a second GLM, we modelled all trials in a block with a single regressor with a parametric modulator constructed from the group-mean RT curve. Apart from the motion covariates, all other regressors were convolved with SPM’s ‘canonical’ hemodynamic response function (hrf) composed of linear combination of two gamma functions. A high-pass filter of 128 s was applied to the predicted and measured timecourses and the data were pre-whitened

GLMs were fit to each voxel’s timecourse for whole-brain analyses and also to the mean time-course of pre-defined regions of interest (ROIs). Three sets of ROIs incorporating the striatum and task control regions in frontal and parietal cortex were analysed. Two ‘striatum’ ROIs were defined as the ‘Caudate’ and ‘Putamen’ regions from the automatic anatomical labelling (AAL) atlas. Six cortical ROIs were identified from the ‘multiple-demand’ (MD) network (Fedorenko et al., 2013) including the anterior PFC (aPFC), dorsolateral PFC (dlPFC), anterior insula (AI), inferior frontal junction (IFJ), dorsal ACC (dACC) and inferior parietal sulcus (IPS). Previous work has suggested that these MD ROIs segregate into separate ‘fronto-parietal’ (DLPFC, IFJ, IPS) and ‘cingulo-opercular’ networks (aPFC, AI, dACC) (Crittenden et al., 2016). Therefore, we treated these as separate networks for our analyses.

We also ran a generalized psychophysical interaction (gPPI) analysis. We defined the seed region as the ‘caudate’ region in the AAL atlas. This caudate region was employed as a mask for extracting the activity timeseries data of the region from the high-pass filtered functional data. These values then served as a regressor (without hrf convolution) and was employed as a regressor along with all other regressors for each trial position, allowing full interactions between the regressors (McLaren et al, 2012). The interaction regressors of the caudate timeseries and each trial position were then subjected to a RT-weighted contrast.

Summary statistics from the individual participant GLM fits were then entered into a 2nd level random-effects analysis to combine data across participants. We evaluated three primary contrasts. First, we contrasted early trials (first 4) against late trials (last 4) in the block, separately for the first and second half of the experiment to search for voxels that show elevated activity on the early trials. Second, we examined the parametric modulation of trial activity with the z-scored group-mean RT curve to identify voxels which showed activity tracking the shape of the mean RT-curve. The early vs late contrast and the gPPI contrast was cluster-corrected to p < 0.05 family wise error (FWE) using Gaussian random field theory. The RT-curve contrast -maps was also corrected with a more conservative FWE height correction at the individual voxel level to lower false positive rates for the sensitive trend analysis contrasts. Height correction was also important for this analysis as cluster correction can be biased against smaller structures, like the striatum, which was a focus of this analysis from our hypotheses.

Cluster peak co-ordinates from all whole-brain contrasts are listed in Tables 2 to 4. All cluster peaks greater than 8 mm apart are listed for cluster-corrected (*p* = 0.05 FEW) statistical maps. Cluster forming height threshold is always at p < 0.001. Region labels are based on the AAL2 atlas. For peak co-ordinates not listed in the AAL2 atlas, the top region association in Neurosynth (Yarkoni et al., 2011) are listed in parentheses.

**Table 2:** fMRI activations - Figure 4*

| Region (AAL2) | No. of voxels | MNI Coordinates |  |  | Peak t-value |
| --- | --- | --- | --- | --- | --- |
|  |  | x | y | z |  |
| Early trials > Late trials - All blocks (k = 356 voxels) |  |  |  |  |  |
| Location not in atlas (thalamus) | 2458 | 6 | -20 | -8 | 7.9588 |
| Left Thalamus |  | -8 | -16 | -2 | 6.1781 |
| Right Pallidum |  | 28 | -12 | -6 | 5.4447 |
| Right Thalamus |  | 12 | -20 | 8 | 5.3053 |
| Right Thalamus |  | 16 | -14 | 14 | 5.1011 |
| Location not in atlas (thalamus) |  | -20 | -14 | 14 | 4.7974 |
| Location not in atlas (thalamus) |  | 24 | -16 | 8 | 4.6239 |
| Location not in atlas (thalamus) |  | -14 | -6 | 12 | 4.5113 |
| Right Caudate |  | 12 | 12 | -8 | 4.4761 |
| Location not in atlas (putamen) |  | 18 | 0 | 10 | 4.4399 |
| Right Pallidum |  | 18 | 2 | -2 | 4.2306 |
| Location not in atlas (thalamus) |  | -8 | -2 | 4 | 4.0822 |
| Location not in atlas (basal ganglia) |  | -20 | 0 | 14 | 4.0396 |
| Right inferior parietal gyrus | 1801 | 52 | -40 | 56 | 6.8505 |
| Right angular gyrus |  | 38 | -72 | 46 | 6.1301 |
| Right middle occipital gyrus |  | 32 | -74 | 36 | 5.7962 |
| Right supramarginal gyrus |  | 62 | -48 | 36 | 5.0075 |
| Location not in atlas (intraparietal sulcus) |  | 28 | -56 | 40 | 4.7586 |
| Right inferior parietal gyrus |  | 36 | -54 | 52 | 4.6694 |
| Right inferior parietal gyrus |  | 32 | -50 | 46 | 4.4428 |
| Right inferior parietal gyrus |  | 42 | -46 | 38 | 4.0727 |
| Right supramarginal gyrus |  | 56 | -42 | 38 | 3.9844 |
| Left inferior parietal gyrus | 1621 | -48 | -60 | 50 | 6.5381 |
| Left inferior parietal gyrus |  | -56 | -46 | 48 | 6.1328 |
| Left inferior parietal gyrus |  | -38 | -62 | 52 | 5.9828 |
| Left inferior parietal gyrus |  | -52 | -46 | 40 | 5.352 |
| Left inferior parietal gyrus |  | -34 | -78 | 40 | 5.1028 |
| Left supplementary motor area | 356 | -6 | 22 | 48 | 6.2597 |
| Right median cingulate & paracingulate gyri |  | 8 | 26 | 38 | 4.2527 |
| Left superior frontal gyrus, medial |  | 2 | 34 | 44 | 3.7722 |
| Right superior frontal gyrus, medial |  | 8 | 26 | 48 | 3.5992 |
| Right Inferior frontal gyrus, opercular part | 2231 | 48 | 10 | 26 | 6.0965 |
| Right precentral gyrus |  | 54 | 14 | 40 | 6.0404 |
| Right middle frontal gyrus |  | 44 | 32 | 22 | 5.5242 |
| Right precentral gyrus |  | 40 | 4 | 30 | 5.4625 |
| Right middle frontal gyrus |  | 44 | 10 | 54 | 5.3391 |
| Right inferior frontal gyrus, triangularis |  | 44 | 18 | 24 | 4.6771 |
| Right inferior frontal gyrus, opercularis |  | 46 | 16 | 6 | 4.5817 |
| Right insula |  | 32 | 22 | -2 | 4.5597 |
| Right middle frontal gyrus |  | 34 | 8 | 62 | 4.1642 |
| Right superior frontal gyrus |  | 26 | 8 | 52 | 4.1347 |
| Right inferior frontal gyrus, triangularis |  | 56 | 22 | 16 | 3.7697 |
| Left superior frontal gyrus | 377 | -24 | 6 | 64 | 5.891 |
| Left middle frontal gyrus |  | -28 | 14 | 58 | 5.8071 |
| Left middle frontal gyrus |  | -40 | 8 | 54 | 4.3065 |
| Left middle frontal gyrus | 406 | -50 | 20 | 38 | 5.028 |
| Left inferior frontal gyrus, triangularis |  | -46 | 30 | 22 | 4.2496 |
| Left middle frontal gyrus |  | -42 | 26 | 32 | 4.1584 |
| Left middle frontal gyrus |  | -44 | 36 | 32 | 4.1286 |
| Left inferior frontal gyrus, triangularis |  | -38 | 18 | 26 | 3.9437 |
| Left middle frontal gyrus |  | -44 | 18 | 48 | 3.8292 |
| Left lingual gyrus | 1049 | -20 | -78 | -10 | -7.1867 |
| Left cuneus |  | -10 | -94 | 26 | -4.8661 |
| Right middle occipital gyrus | 1122 | 28 | -88 | 20 | -5.8161 |
| Right lingual gyrus |  | 22 | -82 | -4 | -5.0473 |

| Early trials > Late trials - blocks 1-10 (k = 298 voxels) |  |  |  |  |  |
| --- | --- | --- | --- | --- | --- |
| Right Inferior frontal gyrus, opercular part | 2990 | 46 | 8 | 24 | 8.5075 |
| Right middle frontal gyrus |  | 44 | 6 | 52 | 5.7576 |
| Right middle frontal gyrus |  | 46 | 36 | 18 | 5.7336 |
| Left middle frontal gyrus | 5652 | -52 | 20 | 36 | 6.896 |
| Left thalamus |  | -12 | -16 | 6 | 6.8357 |
| Location not in atlas (substantia nigra) |  | 6 | -20 | -12 | 6.5943 |
| Left inferior frontal gyrus, opercularis |  | -52 | 10 | 10 | 5.8679 |
| Right pallidum |  | 16 | 2 | 0 | 5.4844 |
| Left middle frontal gyrus |  | -28 | 16 | 56 | 5.463 |
| Left insula |  | -30 | 22 | 0 | 5.3038 |
| Location not in atlas (basal ganglia) |  | 28 | -20 | 4 | 4.2829 |
| Right anterior insula |  | 30 | 26 | 0 | 4.2525 |
| Left putamen |  | -12 | 6 | -10 | 3.8611 |
| Left superior parietal gyrus | 1932 | -36 | -64 | 54 | 6.6421 |
| Left inferior parietal gyrus |  | -52 | -38 | 46 | 5.7287 |
| Left middle occipital gyrus |  | -30 | -74 | 28 | 3.6659 |
| Left inferior parietal gyrus |  | -48 | -50 | 42 | 4.2473 |
| Right inferior parietal gyrus | 1143 | 54 | -40 | 54 | 6.0063 |
| Right angular gyrus |  | 38 | -72 | 46 | 5.2242 |
| Location not in atlas (intraparietal sulcus) |  | 34 | -44 | 38 | 3.6628 |
| Right cuneus | 533 | 20 | -96 | 10 | -5.7556 |
| Left lingual gyrus | 376 | -18 | -82 | -10 | -5.3105 |
| Left superior occipital gyrus |  | -14 | -98 | 14 | -4.244 |

| Early trials > Late trials - blocks 11-20 (k = 271 voxels) |  |  |  |  |  |
| --- | --- | --- | --- | --- | --- |
| Right thalamus | 271 | 4 | -14 | 0 | 5.6013 |
| Right angular gyrus | 497 | 40 | -70 | 40 | 5.2922 |
| Right cuneus | 1751 | 20 | -80 | 26 | -4.0053 |
| Right middle occipital gyrus |  | 30 | -86 | 20 | -6.8302 |
| Left superior occipital gyrus |  | -14 | -84 | 26 | -5.8452 |
| Left lingual gyrus | 513 | -18 | -78 | -10 | -6.7048 |

\*all cluster peaks greater than 8 mm apart for the Early vs Late trial contrasts (cluster-corrected $p = 0.05$ FWE) shown in Figure 4. Cluster forming height threshold is always $p < 0.001$ . Number of voxels refers to the size of the cluster, listed once per cluster. Region labels are based on the AAL2 atlas. For peak co-ordinates not listed in the AAL2 atlas, the top region association in Neurosynth (Yarkoni et al., 2011) is listed in parentheses.

**Table 3:** fMRI activations - Figure 5

| Region (AAL2) | No. of voxels | MNI Coordinates |  |  | Peak t-value |
| --- | --- | --- | --- | --- | --- |
|  |  | x | y | z |  |
| RT parametric > baseline (k = 359) |  |  |  |  |  |
| Right parahippocampal gyrus | 41951 | 18 | -42 | -10 | 7.8862 |
| Right putamen |  | 16 | 12 | -6 | 7.6902 |
| Right precentral gyrus |  | 56 | 12 | 38 | 7.4561 |
| Left putamen |  | -10 | 10 | -10 | 7.147 |
| Right olfactory cortex |  | 8 | 26 | -12 | 7.0611 |
| Left insula |  | -36 | 10 | -12 | 7.0577 |
| Location not in atlas (orbitofrontal cortex) |  | 22 | 18 | -12 | 6.9925 |
| Right middle occipital gyrus |  | 34 | -74 | 38 | 6.7987 |
| Left median cingulate & paracingulate gyri |  | -12 | -22 | 44 | 6.7703 |
| Right middle temporal gyrus |  | 54 | -70 | 8 | 6.7131 |
| Right anterior cingulate and paracingulate gyri |  | 4 | 32 | 14 | 6.5716 |
| Location not in atlas (striatum) |  | -8 | 4 | -2 | 6.5084 |
| Right precentral gyrus |  | 40 | -6 | 62 | 6.3867 |
| Right olfactory cortex |  | 22 | 12 | -18 | 6.3313 |
| Left insula |  | -40 | 0 | -10 | 6.3281 |
| Right insula |  | 38 | 12 | -10 | 6.2657 |
| Right insula |  | 40 | 4 | -6 | 6.2104 |
| Left anterior cingulate and paracingulate gyri |  | -6 | 32 | 16 | 6.2045 |
| Left anterior cingulate and paracingulate gyri |  | -6 | 22 | 26 | 6.1912 |
| Location not in atlas (striatum) |  | -22 | 20 | -8 | 6.1892 |
| Left inferior parietal gyrus |  | -44 | -42 | 48 | 3.7144 |
| Left middle temporal gyrus | 359 | -54 | -72 | 8 | 6.5447 |
| Left middle occipital gyrus |  | -48 | -82 | 2 | 4.6365 |
| Left middle temporal gyrus |  | -62 | -56 | 8 | 4.0767 |
| Left middle frontal gyrus | 911 | -30 | 52 | 28 | 6.4783 |
| Left middle frontal gyrus |  | -40 | 38 | 34 | 6.1806 |
| Left middle frontal gyrus |  | -32 | 42 | 36 | 4.9541 |
| Left middle frontal gyrus, triangularis |  | -46 | 30 | 22 | 4.2401 |
| Left middle frontal gyrus, triangularis |  | -48 | 44 | 4 | 4.1996 |
| Left middle frontal gyrus, triangularis |  | -32 | 30 | 26 | 4.0885 |
| Left middle frontal gyrus |  | -40 | 50 | 18 | 4.0718 |
| Left middle frontal gyrus |  | -28 | 40 | 22 | 4.0117 |

\*cluster peaks greater than 8 mm apart for the group RT parametric > baseline contrast (cluster-corrected $p = 0.05$ FWE) shown in Figure 5. Cluster forming height threshold is $p < 0.001$ .

**Table 4:**
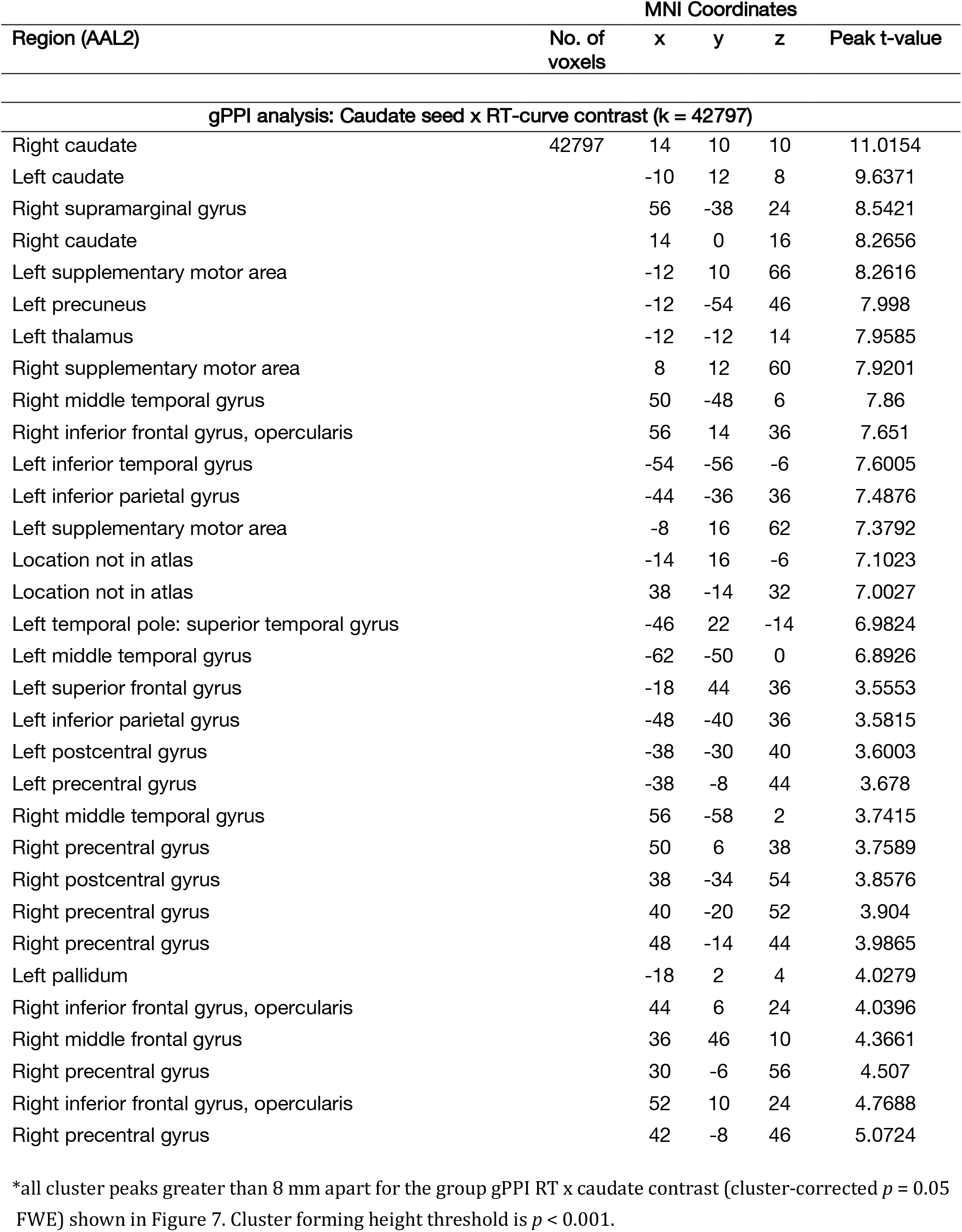
fMRI activations - Figure 7

| Region (AAL2) | No. of voxels | MNI Coordinates |  |  | Peak t-value |
| --- | --- | --- | --- | --- | --- |
|  |  | x | y | z |  |
| gPPI analysis: Caudate seed x RT-curve contrast (k = 42797) |  |  |  |  |  |
| Right caudate | 42797 | 14 | 10 | 10 | 11.0154 |
| Left caudate |  | -10 | 12 | 8 | 9.6371 |
| Right supramarginal gyrus |  | 56 | -38 | 24 | 8.5421 |
| Right caudate |  | 14 | 0 | 16 | 8.2656 |
| Left supplementary motor area |  | -12 | 10 | 66 | 8.2616 |
| Left precuneus |  | -12 | -54 | 46 | 7.998 |
| Left thalamus |  | -12 | -12 | 14 | 7.9585 |
| Right supplementary motor area |  | 8 | 12 | 60 | 7.9201 |
| Right middle temporal gyrus |  | 50 | -48 | 6 | 7.86 |
| Right inferior frontal gyrus, opercularis |  | 56 | 14 | 36 | 7.651 |
| Left inferior temporal gyrus |  | -54 | -56 | -6 | 7.6005 |
| Left inferior parietal gyrus |  | -44 | -36 | 36 | 7.4876 |
| Left supplementary motor area |  | -8 | 16 | 62 | 7.3792 |
| Location not in atlas |  | -14 | 16 | -6 | 7.1023 |
| Location not in atlas |  | 38 | -14 | 32 | 7.0027 |
| Left temporal pole: superior temporal gyrus |  | -46 | 22 | -14 | 6.9824 |
| Left middle temporal gyrus |  | -62 | -50 | 0 | 6.8926 |
| Left superior frontal gyrus |  | -18 | 44 | 36 | 3.5553 |
| Left inferior parietal gyrus |  | -48 | -40 | 36 | 3.5815 |
| Left postcentral gyrus |  | -38 | -30 | 40 | 3.6003 |
| Left precentral gyrus |  | -38 | -8 | 44 | 3.678 |
| Right middle temporal gyrus |  | 56 | -58 | 2 | 3.7415 |
| Right precentral gyrus |  | 50 | 6 | 38 | 3.7589 |
| Right postcentral gyrus |  | 38 | -34 | 54 | 3.8576 |
| Right precentral gyrus |  | 40 | -20 | 52 | 3.904 |
| Right precentral gyrus |  | 48 | -14 | 44 | 3.9865 |
| Left pallidum |  | -18 | 2 | 4 | 4.0279 |
| Right inferior frontal gyrus, opercularis |  | 44 | 6 | 24 | 4.0396 |
| Right middle frontal gyrus |  | 36 | 46 | 10 | 4.3661 |
| Right precentral gyrus |  | 30 | -6 | 56 | 4.507 |
| Right inferior frontal gyrus, opercularis |  | 52 | 10 | 24 | 4.7688 |
| Right precentral gyrus |  | 42 | -8 | 46 | 5.0724 |

\*all cluster peaks greater than 8 mm apart for the group gPPI RT x caudate contrast (cluster-corrected $p = 0.05$ )
FWE) shown in Figure 7. Cluster forming height threshold is $p < 0.001$ .

For ROI analyses, regression weights from the individual participants were evaluated in a random-effects analysis with repeated measures ANOVAs with region and trial number as within-subject factors. A second ANOVA included region, trial number and experiment half as factors. Differences across ROIs in the size of the first trial RT effect was evaluated with a repeated measures ANOVA (detailed in the results section).

## Results

All participants reached criterion on the task with a maximum of 4 rounds of practice on the full task (mode=2). In the final, feedback-free practice block prior to scanning, mean accuracy was 89% and mean RT was 1245 ± 136 ms. In the scanner, participants (not including those excluded) had an overall accuracy of 89.4% ± 3%, and RT of 1439 ± 147 ms.

### Novel sequential trial structures induce slower RTs on the first trial, followed by rapid decline on subsequent trials

To track the implementation of control policies, we examined changes in behavioral measures as a function of the number of trials within a block, as participants gained experience with a novel sequential trial structure. Error rates (Fig 3(a)) did not show any significant change as a function of number of trials (One-way ANOVA: *F*_11,220_ = 1.184, *p* > 0.25, *η_p_^2^* = 0.06). On the other hand, we observed a significant main effect of the number of trials in a block on RT (Oneway ANOVA: *F*_11,220_ = 2.485, *p* = 0.006, *η_p_^2^* = 0.11), which was driven by slower responses on the very first trial of a block (Fig 3(b)). Pair-wise *t*-tests of adjacent trials showed that RTs declined rapidly after the first trial (Paired t-test 1^st^ vs 2^nd^ trial RT: *t*_20_ 3.94, *p_uncorrected_* < 0.001, *p_corrected_* = 0.009, *d* = 0.86. All other pair-wise tests: *p_uncorrected_* > 0.1). There was a significantly larger decline in RT between the first two trials, than between the second and the third (Paired *t*-test Trial 1-2 ΔRT vs Trial 2-3 ΔRT: *t*_20_ =3.47, *p* = 0.002, *d* = 0.76). In other words, blocks with novel trial structures induce a slowing of responses on the first trial, even when task rules are highly familiar. However, participants rapidly adapt to the novel trial structure.

**Figure 3.**
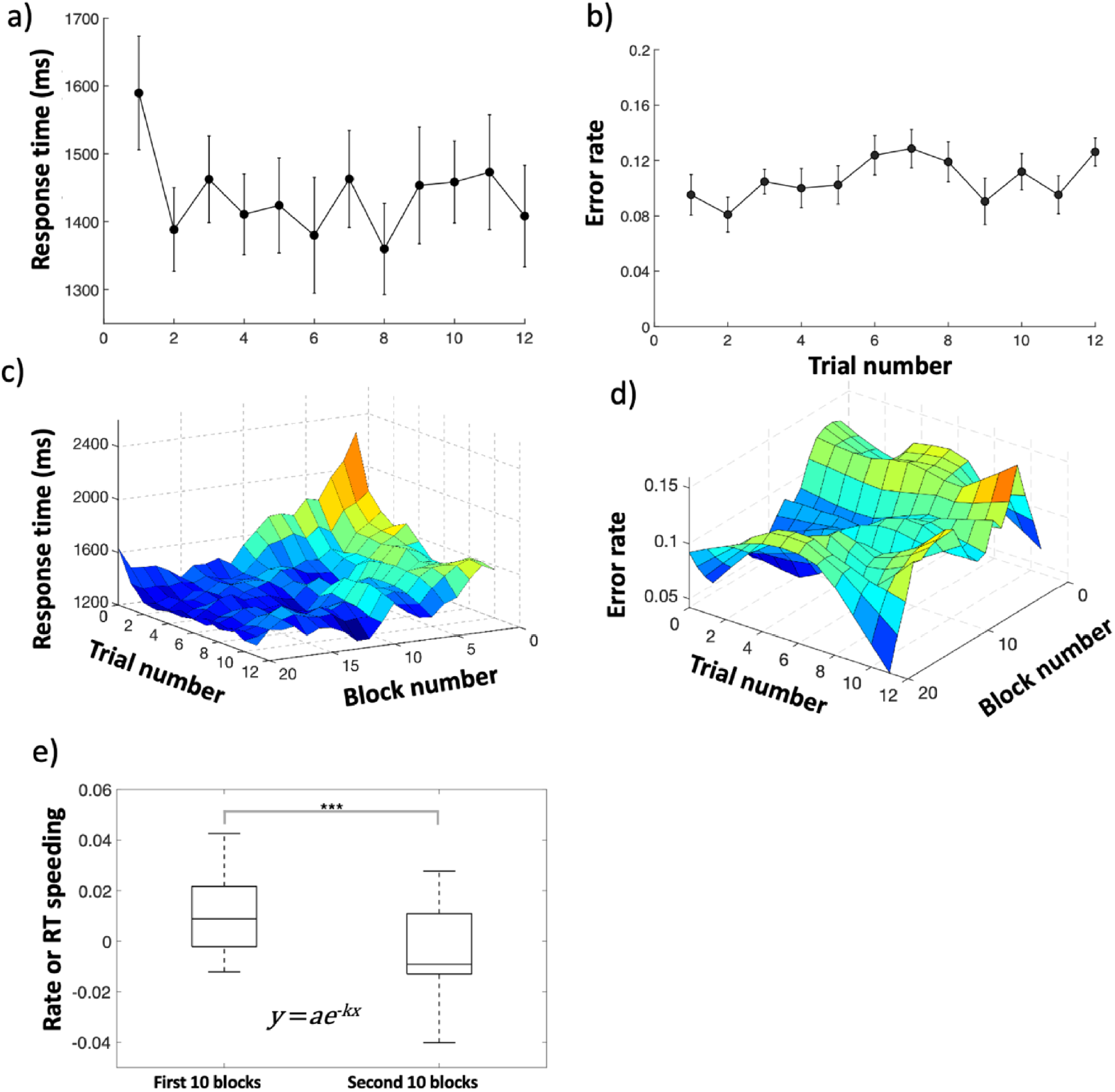
Behavioral results. (a) Plot shows error rates as a function of trial number. Error rates are largely stable, with no significant effect of the number of trials experienced. (b) Plot shows mean RTs as a function of trial number. Participants are slow on the very first trial on encountering a novel trial structure, but rapidly speed their responses to asymptotic levels. (c and d) Mean RTs and error rates as a function of both trial and block numbers. Data were smoothed along both trial and block axes prior to plotting for illustration. As participants experience more novel trial structures, the slow responses on early trials are abolished. (e) This effect is confirmed by a formal comparison of estimated (exponential) rates of speeding in first v/s last 10 blocks.

### First trial slowing induced by novel sequential trial structures is abolished across blocks as more trial structures are experienced

We next examined how these within-block RT dynamics changed across blocks as participants experience an increasing number of novel sequential trial structures. As Figure 3(c) illustrates, we observed a flattening of the RT dynamics across blocks, with the pattern of slower responses on the early trials being abolished after 10 blocks. To formally test this, we fit an exponential decay function to each participant’s smoothed RT curves, separately for the first (blocks 1-10) and second (blocks 11-20) half of the experiment (Fig 3(d)). The rate of decay (k) was significantly higher in the first half vs second half (Paired *t*-test 1^st^ vs 2^nd^ half rate of decay (k): *t*_20_ =3.91, *p* < 0.001, *d* = 0.85). This difference was driven primarily by a significant change in the first trial cost (difference between 1^st^ trial RT and mean RT of trials 2-12), which reduced from 273 ms in the first half of the experiment to a small benefit in the second half (−12 ms) of the experiment (Paired *t*-test 1^st^ vs 2^nd^ half rate of decay (k): *t*_20_ =3.25, *p* = 0.004, *d* = 0.72)

An analogous analysis of error rates did not show any significant differences in the rate of decay (Paired *t*-test 1^st^ vs 2^nd^ half rate of decay (k): *t*_20_ =1.1, *p* > 0.1, *d* = 0.24).

This pattern of results suggests two distinct, interacting processes tied to the implementation of control policies for efficiently performing a task. One process unfolds with the first encounter with a novel sequential structure, even in the context of highly familiar rules, and enables the rapid implementation of an efficient control policy tuned to the novel sequential trial structure over the first few trials. A second process unfolds at a longer time scale as participants encounter multiple different novel sequential trial structures, and enables faster adaptation to novel trial structures. This slower learning process may reflect the emergence of an abstract control policy or consistent strategy that participants can apply with lower variance to a variety of trial structures. Alternatively, it may reflect changes in underlying task representations, that enable more efficient coding of trial cues independent of trial sequences.

### Fronto-parietal regions and striatum show elevated activity on the early trials with a novel sequential trial structure and track respones time dynamics

Previous work studying the implementation of novel instructed rules has consistently identified elevated activity in fronto-parietal cortex on early trials of the task as one of the primary neural correlates of rapid task learning (Ruge & Wolfensteller, 2010; Hartstra et al., 2011; Ruge & Wolfensteller, 2013). To test whether his finding generalizes to the implementation of control policies in the context of familiar rules, we ran a whole-brain analysis testing an early trial effect, contrasting activity on the early (first 4) vs late (last 4) trials in the block (Fig 4, Table 2).

**Figure 4.**
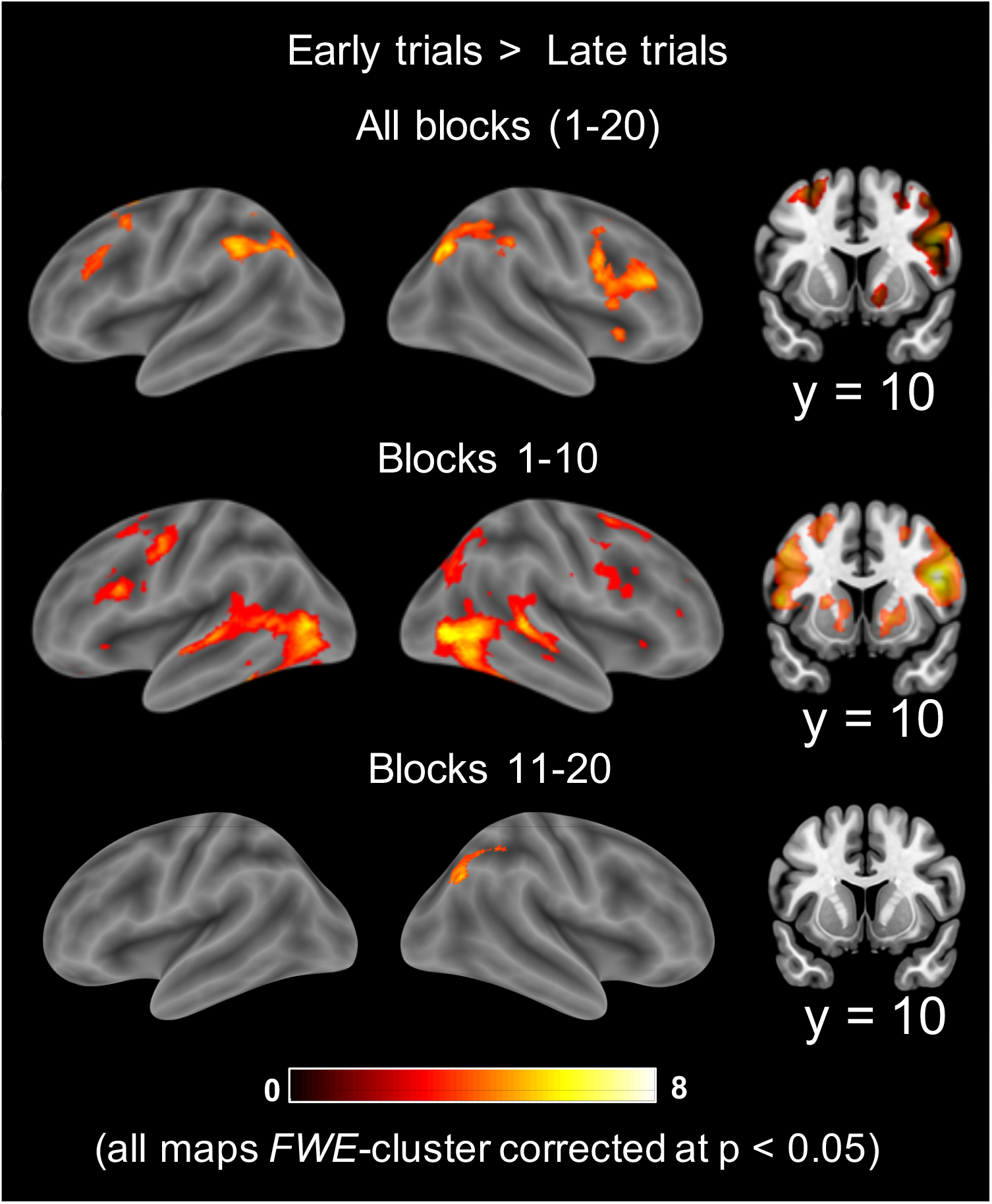
Whole-brain contrast of BOLD activity on first 4 vs last 4 trials in a block where a novel sequential structure is introduced. Top panel shows the contrast for all the blocks. Middle panel shows contrast for the first half of the experiment and bottom panel shows the second half. All contrasts are FWE-cluster-corrected to p <0.05. The left and middle panels show surface renderings. The right panel shows a coronal section at y = 10 exposing the striatum. Contrast identifies a fronto-parietal network as well as the striatum as regions where activity is elevated on early trials. This effect appears to be largely abolished by the second half of the experiment.

Indeed, we observed a broad lateral fronto-parietal network shows elevated activity on early trials in a block. Contrary to previous findings, however, we also find the same early trial effect in the striatum, whereas rule implementation studies have found the reverse effect of higher activity on latter trials (Ruge & Wolfensteller, 2010). Separate contrasts of activity in early vs late trials in the first 10 blocks and last 10 blocks of the experiment (Fig 4 middle and bottom panels) suggest that much of this effect appears in the initial blocks of the experiment. In the first 10 blocks, a broad fronto-parietal network, striatum, as well as regions in the ventral visual stream show an early trial effect. In the last 10 blocks, only a cluster in right inferior parietal sulcus survives multiple comparison correction. However, no clusters survived correction in a direct test of the size of the early trial effect in the first 10 vs last 10 blocks. Note, however, that a formal test of this difference in fMRI is hampered by the long temporal separation between early and late blocks and the use of high pass filtering which removes low frequency signal and noise.

To more closely identify neural activity tracking RT dynamics, we examined the parametric modulation of trial activity with the group-mean RT-by-trial curve in a wholebrain analysis. Fig 5 shows whole-brain surface renderings and a coronal slice from a cluster-corrected map depicting regions where activity dynamics were most consistent with the overall shape of the RT-curve (see also Table To highlight the strongest activations, voxels surviving a more conservative voxel-wise FWE correction are overlaid in pink. While a broad network of regions across the brain track RT, we found the strongest activation in clusters in the anterior insula, striatum and the right lateral PFC.

**Figure 5.**
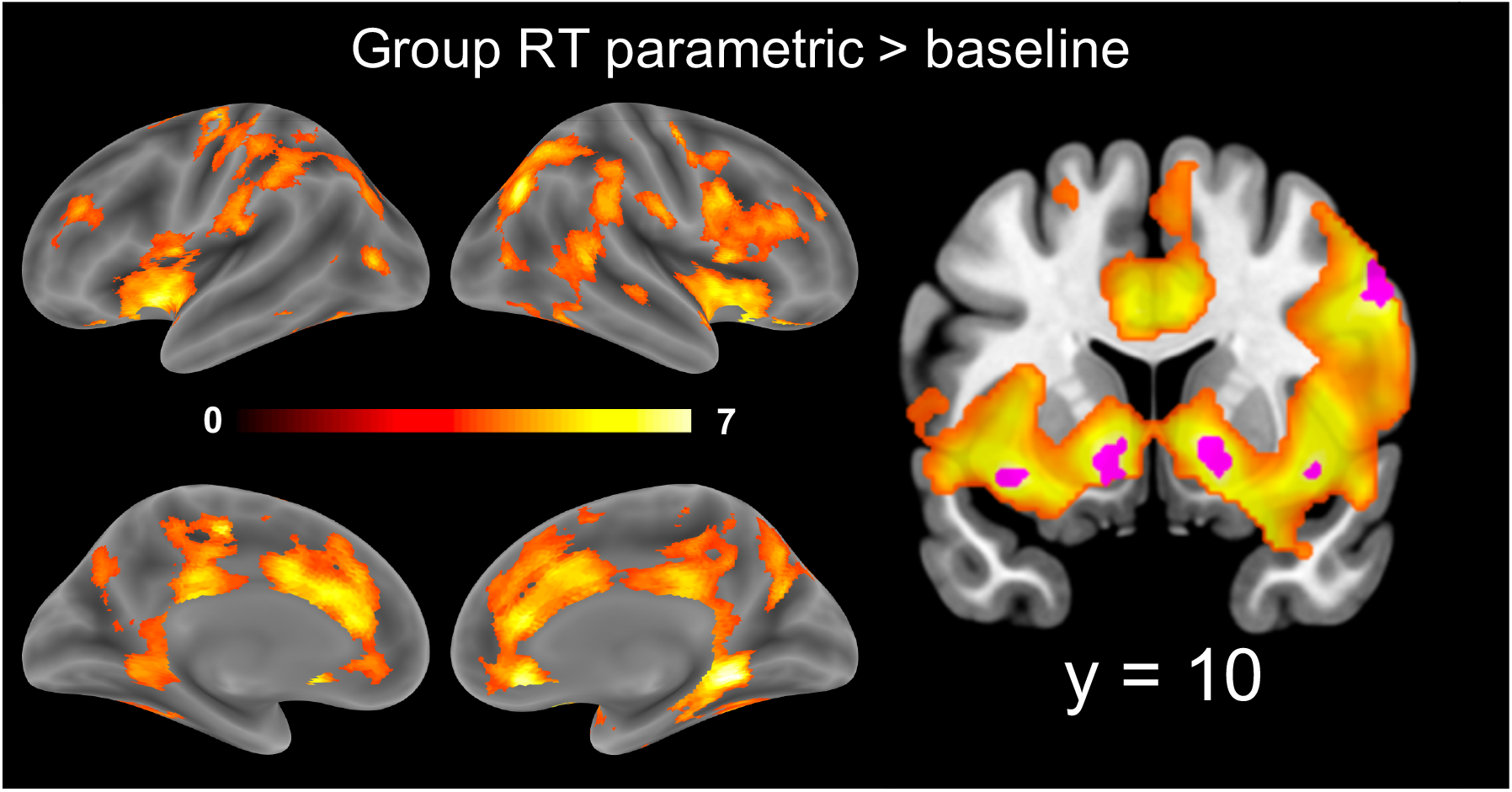
Parametric modulation of trial activity by (z-scored) group-mean RT-by-trial curve. Contrast is cluster-corrected at p < 0.05. Voxels surviving a conservative FWE height correction at the voxel-level are overlaid in pink and represent the strongest activations. Coronal slice (y=10) is depicted showing the strongest activations in the anterior insula, striatum, and right lateral prefrontal cortex (PFC)

### Activity in striatum & the cingulo-opercular network, but not fronto-parietal network shows rapid change

We hypothesized that brain networks associated with cognitive control would be involved in the learning and implementation of control policies. In particular, we focused on ‘multiple-demand’ regions in the cingulo-opercular network (CON), the fronto-parietal network (FPN) as well as the striatum (STR). These regions have been implicated in task control (Dosenbach et al., 2008) and also overlap with regions implicated in rule learning and implementation (Hartstra et al., 2011; Stocco et al., 2012; Muhle-Karbe et al., 2017; Ruge et al., 2019), including of hierarchical rules ((Badre et al., 2010).

We examined the hemodynamic activity dynamics in these regions across trials within a block (Fig 6). Activity dynamics in striatum and the cingulo-opercular network were strikingly similar to within-block RT dynamics, with elevated activity on the first trial followed by a rapid decline (Fig 6(a, b)). On the other hand, activity declined more slowly in the FPN (Fig 6(c). In a 2-way region x trial number rmANOVA, we found significant main effects of trial number (*F*_11,220_ = 8.69, *p* < 0.001, *η_p_^2^* = 0.303) and region (*F*_2,40_ = 13.96, *p* < 0.001, *η_p_^2^* = 0.411), and also, a significant region x trial number interaction (*F*_22,440_ = 2.86, *p* < 0.001, *η_p_^2^* = 0.125). This interaction was driven by differences in the rate of activity decline on early trials across the three regions. Comparing the relative change in activity between trial 1 and 2, versus the change in activity between trial 2 and 3, across the three regions (Fig 6(d)), we found a highly significant region x trial pair interaction (*F*2,40 = 17.25,*p* < 0.001, *η_p_^2^* = 0.463).

**Figure 6.**
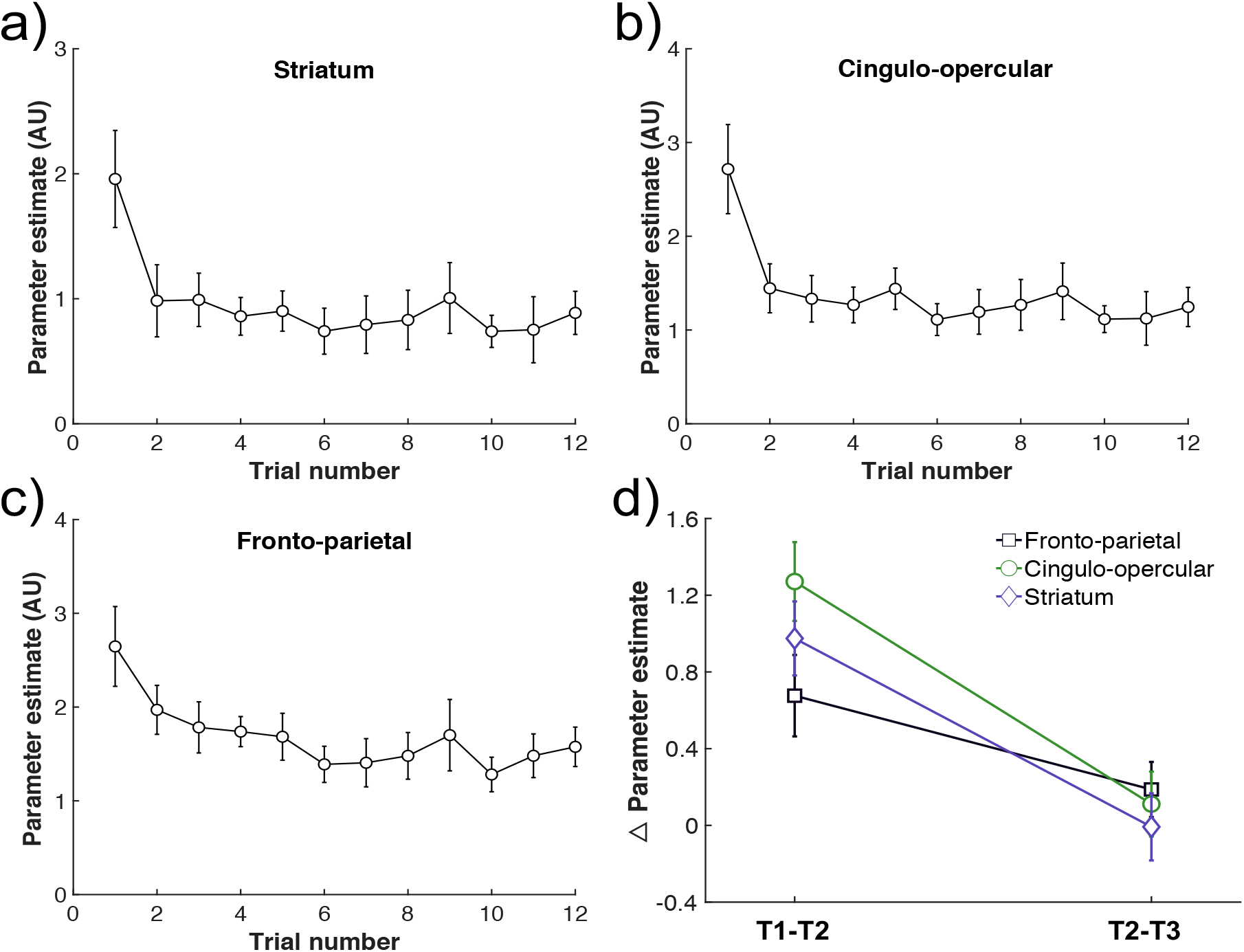
Region-of-interest analysis of trial activity dynamics. Plots (a-c) show trial parameter estimates (regression weights) as a function of trial number for (a) the striatum, (b) multiple-demand (MD) regions in the cingulo-opercular network (dorsal ACC, anterior insula, anterior PFC) and (c) MD regions in the fronto-parietal network (dorsolateral PFC, inferior frontal junction and inferior parietal sulcus). Parameter estimates were averaged bilaterally, and across all ROIs in the network. Separate plots for each ROI are available in the supplementary materials. While all three networks show activity reduction with practice on a novel trial structure, the networks differ on the rate of decrease. (d) Change in parameter estimates from trial 1 to 2 vs trial 2 to 3. Striatum and cingulo-opercular network regions exhibit rapid trial activity dynamics, with a much larger change from trial 1 to 2 compared to trial 2 to 3, with a while frontoparietal network activity reduces at a significantly slower rate (region x trial pair interaction, p < 0.001).

Finally, we asked whether these activity dynamics changed between the first and second half of the experiment. To simplify this analysis, we focused on the relative change in activity between trials 1 and 2 versus 2 and 3. We ran a 3-way region (STR, CON, FPN) x trial pair (1 vs 2, 2 vs 3) x experiment half (1^st^ or 2^nd^). As expected, we found significant main effects of region, (*F*_2,40_ = 9.73, *p* < 0.001, *η_p_^2^* = 0.327) and trial pair (*F*_1,20_ = 10.5, *p* = 0.004, *η_p_^2^* = 0.463), and a significant region x trial pair interaction (*F*_2,40_ = 17.25, *p* < 0.001, *η_p_^2^* = 0.463), but no significant effect of experiment half (*F*_1,20_ = 0.01, *p* > 0.25, *η_p_^2^* <0.001). The region x experiment half, trial pair x experiment half, and the region x trial pair x experiment half interactions were all nonsignificant (all *p* > 0.25). In other words, we found no positive evidence of there being any difference in trial activity dynamics between the first and second half of the experiment.

In summary, we found evidence for rapid activity changes in striatum and the cingulo-opercular network, but not fronto-parietal cortex, which showed a more gradual decline within a block. These results suggest that the fronto-parietal cortex on one hand, and the cingulo-opercular network and striatum on the other, play dissociable roles in control policy learning and implementation. Somewhat surprisingly, despite a significant flattening of RT curves in the second half of the experiment, we found no evidence that trial activity dynamics in either of three regions was different in the second half.

### Striatal functional connectivity with a broad fronto-parietal-temporal network of regions is modulated by RT dynamics

Finally, we asked whether the functional network interactions of the striatum also show dynamics associated with control policy implementation. Novel rule implementation studies by Ruge & colleagues (Ruge & Wolfensteller, 2013; Ruge & Wolfensteller, 2015; Mohr et al., 2016) have shown changes in striatal functional connectivity and integration as a function of practice.

To test whether dynamic interactions can occur as a function of novel control demands, in the absence of novel rule learning or implementation requirements, we ran a generalized psychophysical interaction (gPPI) analysis. We searched for voxels whose activity time courses were differentially corelated with striatal (Caudate) activity timecourses as activity changed in the shape of the group mean RT curve across trials. We found that striatal functional interactions with a broad fronto-parietal-temporal network (Fig 7, Table 4) varied as a function of the RT-curve. In other words, functional interactions of the striatum with this broad network of regions dynamically changed as one experiences a novel trial structure. Finally, we asked the striatum’s interactions with frontoparietal vs cingulo-opercular networks were systematically different, A direct comparison of striatal psychophysical interaction effects with fronto-parietal vs cingulo-opercular networks showed no significant difference (Paired *t*-test FP vs CON: *t*_20_ =1.52, *p* > 0.1, *d* = 0.33).

**Figure 7.**
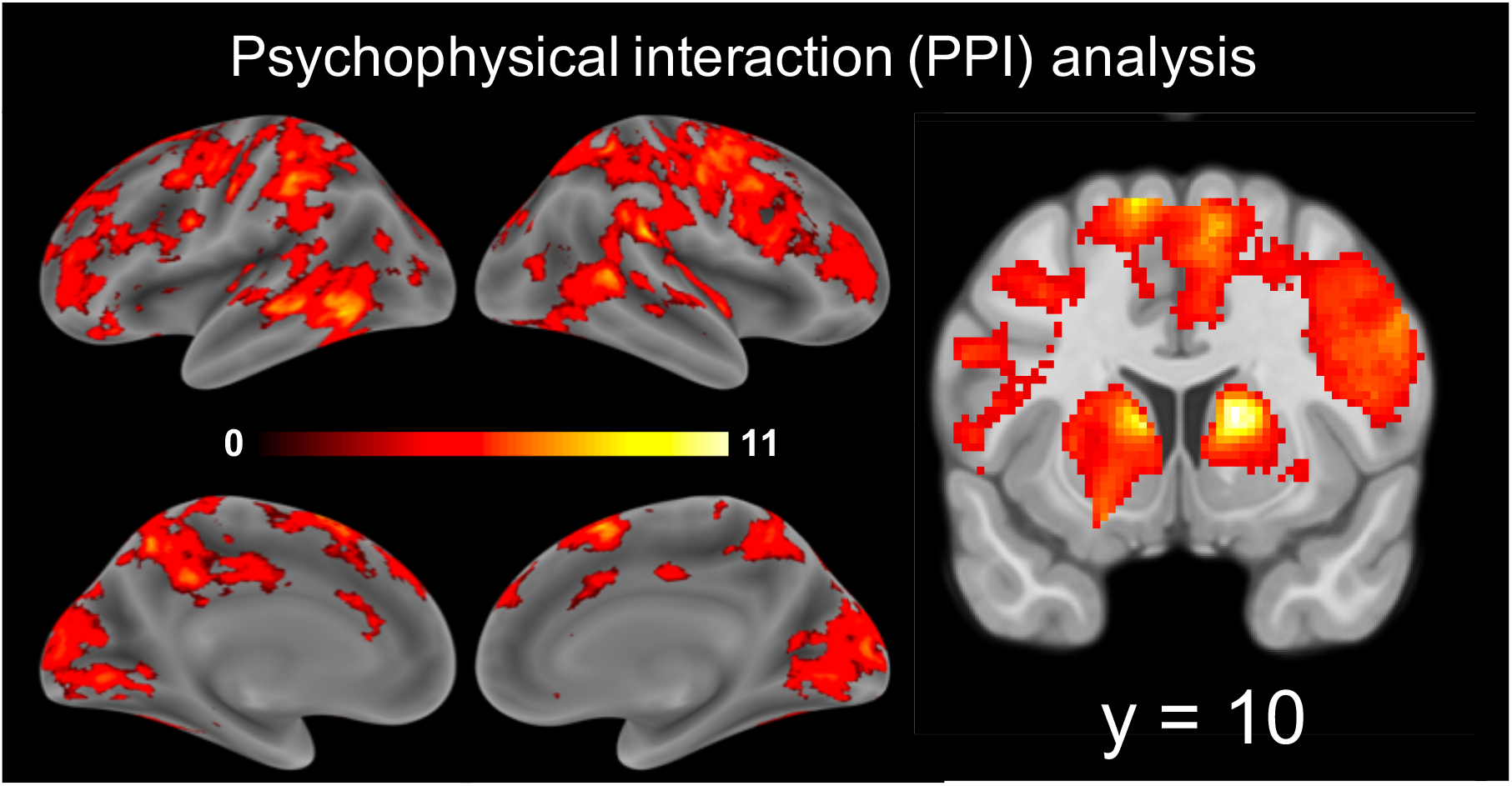
gPPI analysis identified voxels across a broad fronto-parietal-temporal network whose functional connectivity with a seed region in the caudate changes as a function of the RT-curve. Cluster-corrected at p < 0.05. A direct comparison of these striatal-RT interaction effects with fronto-parietal vs cingulo-opercular networks showed no significant difference

## Discussion

Efficient task performance requires implementing a *control policy* that co-ordinates internal cognitive processes to match dynamic task structure. Here, we investigated control policy implementation. We employed a novel 3^rd^ order working memory control task and manipulated the order of cue presentation, thereby altering control demands from block-to-block without changing the stimulus-response (S-R) rules. We find evidence of two adaptive processes, operating over different time scales, that implement efficient, generalizable control policies. We associate these with cortico-striatal control networks.

First, participants were slower to respond on the first trial of a novel task structure, despite the same S-R rules, but sped to asymptote over the next few trials. Hence, implementing a control policy constrains task performance, but implementation is rapid. Note that while first-trial slowing is akin to a ‘switch cost’ (Alport et al., 1994), it is not triggered by a change in S-R rules, but a change in the control demands.

Prior studies have reported similar RT dynamics as evidence of task automatization (Mohr et al., 2016). Our results indicate that this process should be generalized beyond S-R learning to control policy implementation (Bhandari and Badre, 2018). As such, novel S-R rules are not necessary to trigger adaptive processes indexed by early-trial RT dynamics. Rather, S-R rule implementation is a special case of control policy implementation (Bhandari and Badre, 2018) i.e., task model assembly, (Duncan et al., 2008; Bhandari and Duncan, 2014) or control-context learning, (Chiu and Egner, 2019), required when task dynamics change.

Second, the first-trial cost was abolished by the second half of the experiment, with participants exhibiting ‘zero-shot’ adaptation to novel orderings. Such adaptation is remarkable given the complexity of the task, and suggests a second, slower adaptive process as participants experience a variety of task structures.

This slower process might reflect formation of an abstract control policy participants can apply with consistency to a variety of trial structures. For example, consider an output-gating or “waiting” policy, wherein participants store all the cues, postponing all decisions to the end (Chatham et al., 2014; Bhandari and Badre, 2018). This policy uses working memory capacity inefficiently, but can be applied to a variety of task structures consistently.

Changes in task representations themselves may also enable zero-shot adaptation. For example, participants might initially rely on abstract cue representations and employ a multi-step, hierarchical decision process sensitive to the sequential order of the cues. Over time, however, they develop conjunctive representations of different cue combinations, invariant to presentation order. This non-hierarchical conjunctive representation can support a one-step decision robust to order. Future work can distinguish these possibilities.

Third, frontal and parietal regions and the striatum were more active on early trials of a block, especially during the first half of the experiment. Previous fMRI studies of instructed tasks have consistently documented similar activity dynamics in fronto-parietal control networks (Cole et al., 2010; Ruge and Wolfensteller, 2010; Hartstra et al., 2011; Mohr et al., 2016; Hampshire et al., 2019). In these studies, the early engagement of the fronto-parietal network has been associated with encoding of instructed S-R rules. Given that we see similar fronto-parietal dynamics without any change in the S-R rules, we argue that they generalize beyond S-R rules to changes in cognitive control demands.

In one difference from prior work, however, Ruge and Wolfensteller (2010) observed increased striatal activity with practice of novel S-R rules, which we do not observe. Rather, we see elevated striatal activity on the first trial, with subsequent trial activity closely mirroring the group mean RT change. (Ruge and Wolfensteller, 2010) interpreted increasing striatal activity as reflecting the transfer of control to pragmatic rule representations, citing the role of striatum in trial-and-error learning of rules (Boettiger and D’Esposito, 2005; Seger and Cincotta, 2006). The absence of this profile of rising activity in our data may thus reflect the fact that S-R rules do not change in our task.

Importantly, we found that the rate of activity decline dissociated the frontoparietal control network from the cingulo-opercular control network and striatum. All showed greater first-trial activity, but this activity declined more slowly in the frontoparietal cortex. On the other hand, cingulo-opercular regions exhibited a rapid change that more closely matched the changes in RT.

The salience of the first trial might account for the rapid activation change in the cingulo-opercular network, providing an orienting or updating signal to the control system (Menon and Uddin, 2010; Han et al., 2019). The first trial is salient because it is the only trial that alerts participants to the novel control demands. Cingulo-opercular regions have been proposed as a ‘salience network’ (Seeley et al., 2007) and show transient activity at event boundaries (Sridharan et al., 2008). Thus, they may be responding to the salience of these first-trial events. Given that the striatum shows the same fast decline after the first trial, a salience account may similarly apply. Indeed, transient striatal activity has also been observed with salient oddball events (Han et al., 2019), and the caudate is considered a subcortical node of the salience network (Peters et al., 2016).

Alternatively, elevated first-trial activity in striatum and fronto-parietal cortex may reflect gating of control representations in working memory. Several models have proposed cortico-striatal mechanisms for input and output gating of working memory (Frank et al., 2001), and experimental observation has associated gating with a frontal-basal ganglia network (Murty et al., 2011; D’Ardenne et al., 2012; Nee and Brown, 2013; Chatham et al., 2014). On this view, novel control demands trigger gating of control representations in working memory as reflected in cortico-striatal activity. This enables the adaptation observed in behavior. However, fronto-parietal network activity changes lagged behind changes in RTs beyond the first trial. As such, RT dynamics may not reflect processes within fronto-parietal cortex, as has been suggested (Ruge and Wolfensteller, 2010). However, Ruge et al. (2019) recently showed that newly instructed rules are persistently coded in lateral PFC, even though univariate activity falls to baseline levels, suggesting that these activity dynamics possibly index a different process.

Thus, early trial dynamics in striatum and cingulo-opercular activity might be due to salience or to changes in working memory gating policy. Our data cannot distinguish these accounts, and future work should test them. Indeed, they are not mutually exclusive accounts and might separately or jointly characterize the function of these networks.

Finally, functional connectivity between striatum and frontal and parietal regions was modulated by the RT dynamics. Again, this generalized prior observations from S-R rule implementation (Ruge and Wolfensteller, 2010; 2013, 2015; Mohr et al., 2016).

While we did not find reliable evidence for differences in the striatal interactions with cingulo-opercular vs fronto-parietal networks, striatal interactions with different cortical networks may well reflect different functions. We speculate that first-trial connectivity between striatum and cingulo-opercular regions may be driven by information transfer from the cingulo-opercular network to the striatum, perhaps signaling the detection of a salient event and triggering control-related processing. This hypothetical account predicts that first-trial activity in the cingulo-opercular network precedes that in striatum and fronto-parietal cortex. Such latency differences between anterior cingulate cortex and lateral PFC have indeed been observed in humans (Sridharan et al., 2008) and non-human primates (Rothé et al., 2011).

In contrast, striatal-fronto-parietal connectivity may reflect control policy implementation, as predicted by cortico-striatal gating models. However, as already noted, the slow fronto-parietal dynamics lags behavioral adaptation. Notable in this context, Pasupathy and Miller (2005) found learning-related changes in neural activity recorded from caudate nucleus occurred sooner and more abruptly than lateral PFC in a rule-learning study in monkeys, similar to the current study. However, Pasupathy and Miller (2005) found that improved performance correlated with changes in PFC rather than striatum, in contrast to what we find. One possible reconciliation is that learning occurs independently, and at different rates, in striatum and PFC, but behavioral changes may be driven by either or both regions, depending on the task. In the current study, rapid learning in the striatum may be sufficient to support behavioral adaptation to novel orderings.

Finally, while we found extensive neural correlates of rapid adaptation in each block, we could not conclusively identify a neural correlate of the slower adaptation over blocks. In separate contrasts of early-trial effects in each half of the experiment, widespread effects were only detectable in the first half. However, any direct comparison between halves was limited by the necessary removal of low-frequency signal components to filter out low-frequency noise (Friston et al., 1994). Therefore, our power was limited to detect changes across longer time scales. Future work should fill this gap. Indeed, slower adaptation may correlate not only with changes fMRI activity, but also the geometry of representations (Kriegeskorte and Kievit, 2013).

In conclusion, our study identifies two adaptive processes occurring at different time scales that underlie the learning and implementation of efficient, generalizable control policies. We suggest a broader interpretation of activity and connectivity dynamics in a cortico-striatal network than has been previously associated with the learning of S-R rules.

## Acknowledgements

This project was supported by a MURI from the Office of Naval research (N00014-16-1-2832) and by the James S. McDonnell foundation. We thank Celia Ford and Brittany Ciullo for assistance with data collection, and Christopher Chatham & Theresa Desrochers for useful discussions.

## References

Alport A, Styles EA, Hsieh S (1994) 17 Shifting Intentional Set: Exploring the Dynamic Control of Tasks.

Badre D, Kayser AS, D’Esposito MJN (2010) Frontal cortex and the discovery of abstract action rules. 66:315–326.

Bhandari A, Duncan JJC (2014) Goal neglect and knowledge chunking in the construction of novel behaviour. 130:11–30.

Bhandari A, Badre DJC (2018) Learning and transfer of working memory gating policies. 172:89–100.

Boettiger CA, D’Esposito M (2005) Frontal networks for learning and executing arbitrary stimulus-response associations. Journal of Neuroscience 25:2723–2732.

Bourguignon NJ, Braem S, Hartstra E, De Houwer J, Brass M (2018) Encoding of novel verbal instructions for prospective action in the lateral prefrontal cortex: evidence from univariate and multivariate functional magnetic resonance imaging analysis. Journal of cognitive neuroscience 30:1170–1184.

Brass M, Von Cramon DY (2002) The role of the frontal cortex in task preparation. Cerebral cortex 12:908–914.

Brass M, Cramon DYv (2004) Decomposing components of task preparation with functional magnetic resonance imaging. Journal of cognitive neuroscience 16:609–620.

Cao B, Li W, Li F, Li H (2016) Dissociable roles of medial and lateral PFC in rule learning. Brain and Behaviour 6:e00551.

Chatham CH, Frank MJ, Badre D (2014) Corticostriatal output gating during selection from working memory. Neuron 81:930–942.

Chiu Y-C, Egner T (2019) Cortical and subcortical contributions to context-control learning. Neuroscience & Biobehavioral Reviews.

Cole MW, Bagic A, Kass R, Schneider WJJoN (2010) Prefrontal dynamics underlying rapid instructed task learning reverse with practice. 30:14245–14254.

Crittenden BM, Mitchell DJ, Duncan J (2016) Task encoding across the multiple demand cortex is consistent with a frontoparietal and cingulo-opercular dual networks distinction. Journal of Neuroscience 36:6147–6155.

D’Ardenne K, Eshel N, Luka J, Lenartowicz A, Nystrom LE, Cohen JD (2012) Role of prefrontal cortex and the midbrain dopamine system in working memory updating. Proceedings of the National Academy of Sciences 109:19900–19909.

Dosenbach NU, Fair DA, Cohen AL, Schlaggar BL, Petersen SE (2008) A dual-networks architecture of top-down control. Trends in cognitive sciences 12:99–105.

Dumontheil I, Thompson R, Duncan JJJoCN (2011) Assembly and use of new task rules in fronto-parietal cortex. 23:168–182.

Duncan J, Parr A, Woolgar A, Thompson R, Bright P, Cox S, Bishop S, Nimmo-Smith I (2008) Goal neglect and Spearman’s g: competing parts of a complex task. Journal of Experimental Psychology: General 137:131.

Frank MJ, Loughry B, O’Reilly RC (2001) Interactions between frontal cortex and basal ganglia in working memory: a computational model. Cognitive, Affective, & Behavioral Neuroscience 1:137–160.

Friston KJ, Jezzard P, Turner R (1994) Analysis of functional MRI time-series. Human brain mapping 1:153–171.

Hampshire A, Daws RE, Neves ID, Soreq E, Sandrone S, Violante IR (2019) Probing cortical and sub-cortical contributions to instruction-based learning: Regional specialisation and global network dynamics. NeuroImage 192:88–100.

Han SW, Eaton HP, Marois R (2019) Functional fractionation of the Cingulo-opercular network: alerting insula and updating cingulate. Cerebral Cortex 29:2624–2638.

Hartstra E, Kühn S, Verguts T, Brass M (2011) The implementation of verbal instructions: an fMRI study. Human brain mapping 32:1811–1824.

Kriegeskorte N, Kievit RA (2013) Representational geometry: integrating cognition, computation, and the brain. Trends in cognitive sciences 17:401–412.

Menon V, Uddin LQ (2010) Saliency, switching, attention and control: a network model of insula function. Brain Structure and Function 214:655–667.

Miller EK, Cohen JDJAron (2001) An integrative theory of prefrontal cortex function. 24:167–202.

Mohr H, Wolfensteller U, Betzel Rf, Mišić B, Sporns O, Richiardi J, Ruge HJNc (2016) Integration and segregation of large-scale brain networks during short-term task automatization. 7:13217.

Muhle-Karbe PS, Duncan J, De Baene W, Mitchell DJ, Brass M (2017) Neural coding for instruction-based task sets in human frontoparietal and visual cortex. Cerebral Cortex 27:1891–1905.

Murty VP, Sambataro F, Radulescu E, Altamura M, Iudicello J, Zoltick B, Weinberger DR, Goldberg TE, Mattay VS (2011) Selective updating of working memory content modulates meso-cortico-striatal activity. Neuroimage 57:1264–1272.

Nee DE, Brown JW (2013) Dissociable frontal–striatal and frontal–parietal networks involved in updating hierarchical contexts in working memory. Cerebral cortex 23:2146–2158.

Norman DA, Shallice T (1986) Attention to action. In: Consciousness and self-regulation, pp 1–18: Springer.

Pasupathy A, Miller EK (2005) Different time courses of learning-related activity in the prefrontal cortex and striatum. Nature 433:873–876.

Peters SK, Dunlop K, Downar J (2016) Cortico-striatal-thalamic loop circuits of the salience network: a central pathway in psychiatric disease and treatment. Frontiers in systems neuroscience 10:104.

Rothé M, Quilodran R, Sallet J, Procyk E (2011) Coordination of high gamma activity in anterior cingulate and lateral prefrontal cortical areas during adaptation. Journal of Neuroscience 31:11110–11117.

Ruge H, Wolfensteller UJCC (2010) Rapid formation of pragmatic rule representations in the human brain during instruction-based learning. 20:1656–1667.

Ruge H, Wolfensteller U (2013) Functional integration processes underlying the instruction-based learning of novel goal-directed behaviors. NeuroImage 68:162–172.

Ruge H, Wolfensteller U (2015) Distinct fronto-striatal couplings reveal the double-faced nature of response–outcome relations in instruction-based learning. Cognitive, Affective, & Behavioral Neuroscience 15:349–364.

Ruge H, Schäfer TA, Zwosta K, Mohr H, Wolfensteller U (2019) Neural representation of newly instructed rule identities during early implementation trials. eLife 8.

Schneider W, Shiffrin RMJPr (1977) Controlled and automatic human information processing: I. Detection, search, and attention. 84:1.

Seeley WW, Menon V, Schatzberg AF, Keller J, Glover GH, Kenna H, Reiss AL, Greicius MD (2007) Dissociable intrinsic connectivity networks for salience processing and executive control. Journal of Neuroscience 27:2349–2356.

Seger CA, Cincotta CM (2006) Dynamics of frontal, striatal, and hippocampal systems during rule learning. Cerebral cortex 16:1546–1555.

Sridharan D, Levitin DJ, Menon V (2008) A critical role for the right fronto-insular cortex in switching between central-executive and default-mode networks. Proceedings of the National Academy of Sciences 105:12569–12574.

Stocco A, Lebiere C, O’Reilly RC, Anderson JR (2012) Distinct contributions of the caudate nucleus, rostral prefrontal cortex, and parietal cortex to the execution of instructed tasks. Cognitive, affective, & behavioral neuroscience 12:611–628.

Wolfensteller U, Ruge HJFip (2012) Frontostriatal mechanisms in instruction-based learning as a hallmark of flexible goal-directed behavior. 3:192.

Yarkoni T, Poldrack RA, Nichols TE, Van Essen DC, Wager TD (2011) Large-scale automated synthesis of human functional neuroimaging data. Nature methods 8:665.

